# Transcriptional Profiling of Planarian Regeneration Habituating to Physiological Stressor Reveals Individual and Collective Dynamics

**DOI:** 10.64898/2026.07.28.741268

**Authors:** Stefania E. Kapsetaki, Tomer Landsberger, Michael Levin

## Abstract

Exposure to the potassium channel blocker barium chloride (BaCl₂) causes head degeneration in *Dugesia japonica* flatworms, followed by regeneration of BaCl_2_-insensitive heads, offering a unique model for studying transcriptional resilience to novel stress. We performed RNA sequencing on individual planaria to investigate different transcriptional solutions to the BaCl_2_ challenge, and how regeneration history and social environment shape transcriptomic responses to BaCl₂. We identified a robust transcriptional strategy and a potential sub-strategy for enabling BaCl_2_-insensitive head formation. Moreover, we observed pronounced transcriptional differences between untreated worms regenerating from tail fission fragments (tail-regenerated), and untreated full-sized worms that did not fission during the experiment (intact controls), highlighting the lasting impact of regeneration history. Relative to controls, tail-regenerated worms upregulated neurodevelopmental and morphogenetic programs, while downregulating mitochondrial transport and stress-response pathways. Relative to intact controls, BaCl₂-exposed regenerates upregulated ion transport, metabolic, cell cycle, and inflammatory pathways, while downregulating neuronal signaling, ion homeostasis, morphogenesis, and tissue repair programs. Comparison of BaCl₂-exposed isolated and BaCl₂-exposed group-housed worms revealed minimal transcriptional divergence between social conditions. These findings underscore the complex interplay between regeneration, chemical stress, and social context in shaping gene expression.

## Introduction

Throughout history, organisms have faced inevitable stressors that challenge their survival. Resilience — the ability to recover from damage—is a key adaptive trait that shaped the evolutionary trajectories of life. How do cells and tissues determine which of immensely many possible physiological, transcriptional, behavioral, and other actions would enable adaptive response? Understanding this capacity is not only important for evolutionary biology and the study of problem-solving in pre-neural contexts [1, 2], but also a potentially rich source of inspiration for bio-inspired AI, bioengineering, and regenerative medicine approaches [3].

We are interested in how living systems select responses in the space of transcriptional possibilities to solve specific physiological problems – a difficult inverse problem [4, 5]. Barium chloride (BaCl₂), a nonspecific potassium channel blocker, disrupts potassium ion flow and downstream cellular processes [6], which are highly disruptive to tissue because of the importance of potassium flux to neurophysiology and cellular bioelectric signaling more generally [7–10]. We used this reagent as an opportunity to study how living systems adapt to novel challenges.

Planaria are an important model system for studies of responses to severe perturbations, as they are highly regenerative and widely used as model systems for toxicology, stem cell biology, and neuroscience [11–36]. In prior work [6], we showed that planaria experience acute loss of their heads when exposed to BaCl_2_, but remarkably, regenerate BaCl_2_-insensitive heads. We studied the transcriptomic changes of this BaCl_2_ adaptation and found a relatively small number of differentially-expressed genes, including upregulation of TRPMa channels, dampening of cellular communication, and other changes that restore electrical balance offering a unique model for studying transcriptional resilience [37–41]. However, in that study our RNAseq data were generated from pooled worms, revealing a consensus response but leaving open the question of whether individual animals found different (unique) transcriptional solutions to the BaCl_2_ challenge.

Moreover, given recent work on inter-individual assistance in morphogenesis of embryos challenged by teratogens [42], we were interested in possible group dynamics of the response in planaria. Recent [43] and classical data (reviewed in [44]) revealed that large groups can be more resilient to stressors than small groups or individuals due to a fascinating horizontal interaction between conspecifics. In many species, group living buffers individuals against environmental stressors [45]. For example, clustering behavior in *D. dorotocephala* provides protection from ultraviolet radiation [46]. Planaria are also known to exhibit positive thigmotaxis [47], and it is unknown whether the presence of other animals solving the same problem could provide some sort of collective dynamic that alters the course of individual navigation of the transcriptional landscape [48–51]. Would planaria find the same transcriptional responses to regeneration in BaCl_2_ when they are alone versus in a group? We thus compared the transcriptomes of isolated versus group-living regenerating planaria.

Understanding individual variability in response to challenges is critical. Studies in humans and other animals show that stress alters gene expression, but population-level averages obscure individual differences [6, 52]. Some individuals may be more resilient, or may achieve resilience via distinct transcriptional pathways; for example, when rats are exposed to severe stress, specifically fox urine, individuals vary in their transcriptional response to the stressor [53]. Thus, characterization of resilience of a population based on averages is insufficient. In the face of major morphogenetic and physiological disruption, there may be huge diversity in the response across the population. Identifying diversity of successful responses within a population can reveal algorithms by which living material navigates high-dimensional, difficult problem spaces, which in turn can inform personalized approaches in regenerative medicine and bioelectric modulation [54, 55], as well as driving bio-inspired algorithms for solving challenging problems [56, 57].

When planaria are exposed to gap junction blockers, two fundamentally different morphogenetic responses are found in each cohort [58]. Moreover, regeneration of BaCl₂-insensitive heads in planaria provides an opportunity to understand individual variability as well as to control for regeneration-related transcriptional changes – another important aspect of understanding transcriptional responses to physiological challenges, and the specificity of solutions versus general stress response. Thus, we performed individual-level RNA sequencing in a number of conditions to address these open questions. In this study, we investigate: (1) transcriptional diversity among individual planaria with and without BaCl₂ exposure; (2) differences in gene expression between non-BaCl_2_-exposed worms that recently developed from tail pieces vs. long-established individuals; (3) degree to which planaria converge on a common transcriptional solution to head regeneration in BaCl₂; and (4) effects of isolation versus group living on transcriptional adaptation to BaCl₂.

## Methods

### Planaria maintenance

Planaria were maintained following Emmons-Bell’s et al. [6] planaria maintenance protocol. Specifically, we maintained clonal *D. japonica* in dark incubators in Poland Spring water at 13°C. We fed the planaria liver paste once a week, cleaned the colonies approximately two to three hours after feeding, and approximately three days after feeding. In the experiments, we used planaria that we did not feed for seven days and were approximately 0.8-1 cm in length.

### BaCl_2_ experiments

We used two experimental setups to assess the effects of BaCl₂ exposure on: (1) isolated worms (Figure 1a) and (2) isolated versus group-living worms (Figure S2a). We placed isolated worms individually in dishes containing 30 mL of 1 mM BaCl₂, while groups of 30 worms together under identical conditions. We maintained all dishes in the dark at 13°C. In all conditions, we only collected fully regenerated worms for RNA extraction.

**Figure 1.**
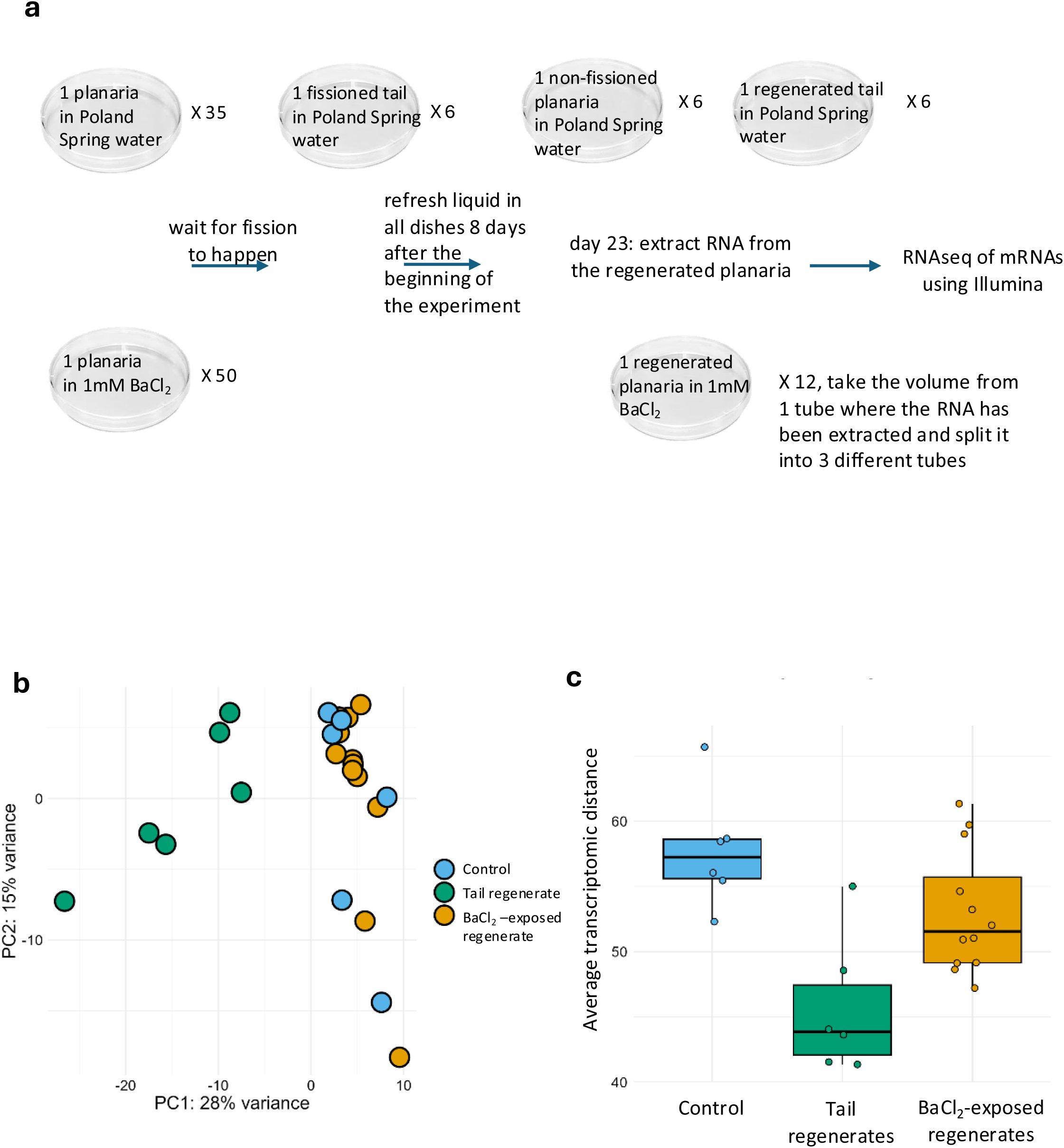
The worms from the three experimental conditions vary in their gene expression. a. Design of experiment testing for transcriptomic differences between isolated versus group-living planaria with regenerated heads in BaCl_2_. b. Principal component analysis (PCA) plot based on normalized gene expression data from planarian samples. The first principal component (PC1) explains 28% of the variance, and the second component (PC2) explains 15% of the variance. Each point represents a biological replicate, colored by experimental condition. Control (blue): non-BaCl_2_-exposed isolated worms. Tail regenerate (green): non-BaCl_2_-exposed worms that recently developed from tail pieces. BaCl_2_ resistant (red): BaCl_2_-exposed isolated regenerates. c. Per-sample average transcriptomic distance among the different conditions. There is higher within-group variability in gene expression in BaCl_2_-resistant worms compared to non-BaCl_2_-exposed worms that recently developed from tail pieces (Dunn’s test, adjusted p-value=0.043), but not in comparison to controls (adjusted p-value=0.13).

### Isolated worms with and without BaCl_2_

We transferred a worm in 30 mL of Poland Spring water in a non-cell-culture-treated dish (size: 100 x 15 mm) in the dark at 13°C (35 replicates). We also transferred a worm in 30 mL of 1 mM BaCl_2_ in Poland Spring water in a non-cell-culture-treated dish (size: 100 x 15 mm) in the dark at 13°C (50 replicates). Some planaria divided in both conditions, and some planaria in BaCl_2_ died during the experiment. The majority of planaria in BaCl_2_ lost their heads and then regenerated their heads. We replaced the liquid in the dishes 8 days after the beginning of the experiment. When a planaria from the condition with Poland Spring water had fissioned, we removed the tail piece and placed it in a non-cell-culture-treated dish (size: 100 x 15 mm) in the dark with 30 mL Poland Spring water. After 23 days, we collected six non-fissioned worms that were growing in Poland Spring water, six regenerated worms that were from tail pieces growing in Poland Spring water, and 12 non-fissioned worms that had regenerated their head in BaCl_2_. We placed one planaria per Eppendorf tube in preparation for the RNA extraction (Figure 1a).

### Isolated versus group-living worms in BaCl_2_

We exposed planaria living on their own to 30 mL of 1 mM BaCl_2_ in Poland Spring water (10 replicates) in non-cell-culture-treated dishes (size: 100 x 15 mm) in the dark at 13°C. We also exposed groups of planaria (30 planaria per group) to 30 mL of 1 mM BaCl_2_ in Poland Spring water (three replicates) in non-cell-culture-treated dishes (size: 100 x 15 mm) in the dark at 13°C (Figure S2). The planaria lost their heads and some died during the experiment. We replaced the liquid in the dishes every day. After 16 days, we collected each of the fully regenerated worms and placed one planaria per tube in preparation for RNA extraction. We refer to fully regenerated worms as worms that have an indistinguishable phenotype from the non-exposed full-body worms. We extracted RNA from three different worms from the isolated replicates, and extracted RNA from four different worms from each of the group conditions (four worms per dish and extracted RNA from each worm separately).

### RNA extraction

We extracted RNA from each worm based on this protocol (https://www.mrcgene.com/wp-content/uploads/mrc-prot/TRI-Reagent.pdf). Below we provide specifics about each step. We added 100 μL TRI reagent in each Eppendorf (Bio Plas Pestle in 1.5mL G-Tube™ Blue/Natural) that had a planaria. We homogenized each planaria using a pestle. We then added 400 μL TRI reagent in each Eppendorf and waited approximately 30 min. Then, we added 50 μL BCP in each Eppendorf. We shook the Eppendorfs for 30 seconds according to the above protocol. We left the Eppendorf tubes for approximately 10 min at room temperature. We centrifuged the Eppendorf tubes at 12,000 g for 15 min at 4°C. Then, we transferred the aqueous solution from each Eppendorf to a fresh Eppendorf. We added 250 μL isopropanol in each new Eppendorf. Then, we left the samples at room temperature for around 10 min. Then, we centrifuged the Eppendorf tubes at 12,000 g for 8 min at 4°C. Then, we removed the supernatant. We added 75% ethanol (i.e., 125 μL nuclease-free water and 375 μL ethanol) in each Eppendorf. We centrifuged the Eppendorf tubes at 7,500 g for 5 min at 4°C. Then, we removed the ethanol solution and briefly air-dried the Eppendorf tubes for around 5 min. We resuspended the RNA pellets in 10 μL nuclease-free water, incubated the samples for 5 min at 1,400 rpm at 34°C, and tracked their A260/A280 ratios using a Nanodrop spectrophotometer. For one experiment, RNA from a single BaCl₂-treated worm was split into three tubes for technical replication.

### RNA-Seq Data Analysis

#### Quality Control and Preprocessing

We performed quality control of raw single-end RNA-seq reads using FastQC to assess the quality of the sequencing data. Following QC, we trimmed adapter sequences and low-quality bases using Trimmomatic with the following parameters: ILLUMINACLIP:TruSeq3-SE.fa:2:30:10 LEADING:3 TRAILING:3 SLIDINGWINDOW:4:15 MINLEN:36. We then subjected the trimmed reads to a second round of quality assessment using FastQC. This ensured the effectiveness of the trimming process.

#### Read Alignment and Quantification

We pseudo-aligned the trimmed reads to the *D. japonica* dd_Djap_v4 transcriptome reference (Rink lab) from PlanMine v3.0 using Kallisto with default parameters. We used Kallisto to estimate transcript abundances.

#### Functional Annotation with eggNOG and BLASTx

We performed functional annotation of the transcriptome using eggNOG-mapper. We annotated the translated coding sequences (CDS) from the dd_Djap_v4 transcriptome to obtain homologous genes, functional descriptions, gene ontology (GO) terms, and pathway information. Gene names throughout the text refer to annotations assigned via eggNOG-mapper, based on best-matching orthologs from model organisms. Additionally, we performed BLASTx searches against the Swiss-Prot database to identify homologous genes and obtain functional information. We integrated the resulting annotations into the tx2gene mapping to enhance the gene-level data.

#### Transcript-to-Gene Mapping

We converted transcript counts to gene counts using the tximport library in R. We prepared a custom tx2gene mapping file containing transcript IDs and corresponding gene symbols. For entries where gene symbols were missing, we used the gene ID as the identifier. We used the tximport function with the parameters ignoreTxVersion=TRUE and countsFromAbundance=“no”.

#### Differential Expression Analysis

We conducted differential expression analysis using DESeq2, constructed a DESeqDataSet object from the count matrix, and summed the technical replicates. We normalized the counts using DESeq2’s method to account for differences in sequencing depth. We performed the Wald test for differential expression analysis with default parameters. We shrunk the log2 fold changes using the “apeglm” method to moderate large-fold changes from lowly expressed genes and to stabilize variance. We considered genes with an adjusted p-value (FDR) < 0.05 and an absolute log2 fold change > log2(1.5) as statistically significantly differentially expressed.

#### Gene Set Enrichment Analysis (GSEA)

We performed a Gene Set Enrichment Analysis (GSEA) using the clusterProfiler R package on the log2 fold change values from the differential expression analysis. The analysis utilized the Gene Ontology (GO) Biological Process (BP) database (org.Hs.eg.db) with the following parameters: minimum gene set size of 20, maximum gene set size of 500.

We performed GSEA using the topGO package in R, leveraging Gene Ontology (GO) annotations derived from eggNOG. The GO terms assigned by eggNOG to each gene in the dd_Djap_v4 transcriptome provided the basis for the enrichment analysis. For each gene set, we evaluated the statistically significant enrichment in specific GO terms to identify overrepresented biological processes, molecular functions, and cellular components associated with the experimental conditions.

We created a custom topGOdata object with the Biological Process (BP) ontology, and defined a list of statistically significant genes based on differential expression results. We used Fisher’s exact test, along with the “classic” algorithm in topGO, to determine enrichment, allowing the identification of GO terms that are statistically significantly associated with the gene set. The analysis incorporated a hierarchical structure of GO terms, capturing both specific and broad biological functions. We further processed the results to reduce redundancy and highlight key GO terms, providing a clearer picture of the main biological themes represented in the data.

#### Visualization

We generated plots with the RNAseq results using the ggplot2, EnhancedVolcano, and pheatmap R libraries.

## Results

### Recent regeneration drives stronger global transcriptional separation than BaCl₂ exposure

We investigated how BaCl₂ exposure and regeneration from tail fragments shape global transcriptional profiles and inter-individual expression variability. We tested three groups: control non-BaCl₂-exposed intact (non-fissioned nor perturbed) worms (henceforward controls), non-BaCl₂-exposed worms recently regenerated from tail fragments of spontaneous fissioning events (henceforward tails-regenerated), and BaCl₂-exposed worms that regenerated their heads (henceforward BaCl_2_-exposed). Read counts varied within and between these conditions (Figure S1). Principal component analysis (PCA) of normalized RNA-seq data revealed two distinct transcriptomic clusters corresponding to the tails-regenerated worms and to control worms mixed with BaCl_2_-exposed worms (Figure 1b). The separation is mainly along the first principal component (PC1, 28% of variance), while the second component (PC2, 15% of variance) does not clearly separate BaCl_2_-exposed worms from controls. Importantly, this separation suggests that recent regenerative history has a stronger effect on global gene expression than exposure to BaCl₂ alone but that barium chloride exposure does induce a distinct effect.

### Increased within-group transcriptomic variability in controls and BaCl₂-exposed worms compared to tail-regenerated worms

Next, we examined intra-group transcriptional variation. We hypothesized that regeneration of BaCl₂-adapted worms may proceed through divergent transcriptional programs, resulting in increased within-group variability compared to controls. To test this, we quantified transcriptomic dissimilarity by calculating, for each worm, the average expression distance to all others in its group (Figure 1b). A Kruskal–Wallis test revealed significant differences in within-group transcriptomic distances across groups (p=0.005). Post-hoc Dunn’s tests showed that both intact controls and BaCl₂-exposed worms exhibited greater variability than tail-regenerated worms (adjusted p=0.043 for BaCl₂-exposed vs tail-regenerated). However, the difference in interindividual transcriptional variability between intact controls and BaCl₂-exposed worms was not statistically significant (adjusted p=0.13). This suggests that no recent regeneration and regeneration under ionic stress may involve more transcriptomic inter-individual variability than canonical regeneration from a tail piece to a full-sized worm.

### Anatomical recovery does not erase the transcriptome of recent regeneration

Barium-insensitive heads arise via regeneration. Thus, in order to individually assess the contributions of regeneration per se, we first compared the transcriptomes of tails-regenerated worms to controls (which had fissioned in the more distant past). PCA clearly separates the groups (Figure 2a). Differential expression analysis between tails-regenerated and controls identified significant transcriptional divergence (Figure 2b). Out of 39,708 genes analyzed, 647 (1.6%) were significantly upregulated (s-value < 0.05, log₂ fold change [LFC] > 0.41) and 1,206 (3%) were significantly downregulated (Figure 2b). Among the strongly upregulated genes, a homolog of human SPTBN5, encoding spectrin repeats (LFC=8.97), indicated major cytoskeletal reorganization potentially linked to morphogenetic processes. Isoforms annotated as homologs of NUDT18, involved in oxidative DNA damage repair, were also substantially elevated (LFC=6.70 and 6.18), suggesting enhanced management of oxidative stress during regeneration. Additionally, increased expression of the *D. japonica* homolog of KIAA1279 (LFC=5.35), a KIF1-binding protein, suggested altered intracellular trafficking associated with neuronal regeneration and structural reorganization. Conversely, genes strongly downregulated in tails-regenerated included genes annotated as homologs of FAM115C (peptidase M60 family, LFC range: –3.11 to –3.85), potentially reflecting reduced extracellular matrix remodeling or altered proteolytic activity. Other highly downregulated genes, such as PI15 (CRISP family, LFC=–3.82) and SLC47A2 (drug-proton antiporter, LFC=–3.10), further indicated shifts in secretion and membrane transport dynamics. Notably, these transcriptional differences persist more than 10 days after the onset of regeneration, indicating that recent regenerative history leaves molecular signatures despite apparent anatomical recovery. Functional enrichment analysis supported these interpretations, revealing enrichment of developmental processes such as morphogenesis, neuron projection guidance, and visual system development among upregulated genes, while metabolic and stress-response functions characterized genes upregulated in control worms (Figure 2c).

**Figure 2.**
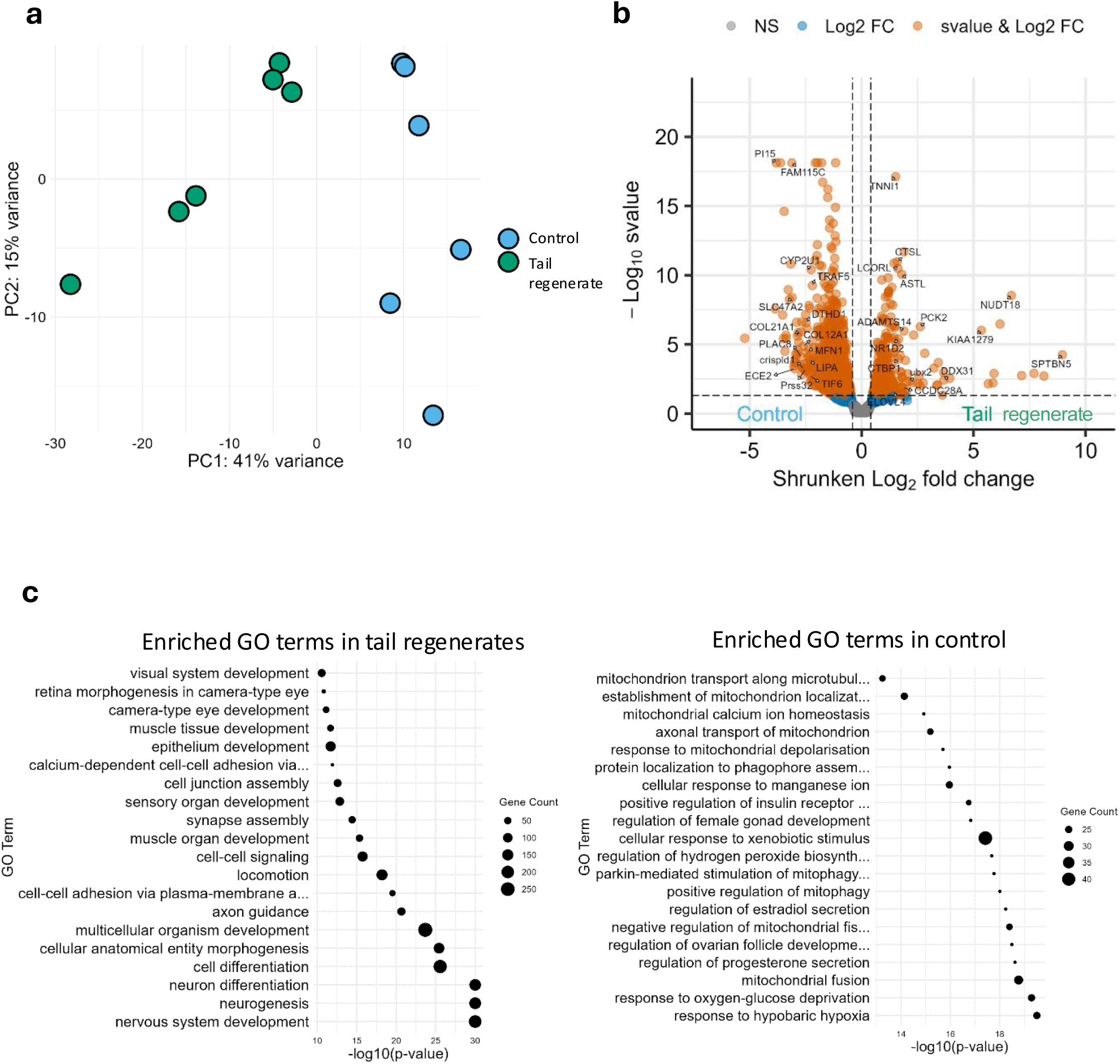
Transcriptional variation among non-BaCl_2_-exposed isolated worms (Control) and non-BaCl_2_-exposed worms that recently developed from tail pieces (Tail regenerate). a. PCA plot based on normalized gene expression data from planarian samples. Each point represents a biological replicate, colored by experimental condition. Control (blue): non-BaCl_2_-exposed isolated worms. Tail regenerate (green): non-BaCl_2_-exposed worms that recently developed from tail pieces. b. 1,853 genes are differentially expressed in non-BaCl_2_-exposed isolated worms (Control) versus non-BaCl_2_-exposed worms that recently developed from tail pieces (Tail regenerate). The volcano plot shows that more genes are upregulated (left) than downregulated (right) in non-BaCl_2_-exposed isolated worms (Control) versus non-BaCl_2_-exposed worms that recently developed from tail pieces (Tail regenerate). More genes are downregulated (left) than upregulated (right) in non-BaCl_2_-exposed worms that recently developed from tail pieces (Tail regenerate) versus non-BaCl_2_-exposed isolated worms (Control). The x-axis represents the shrunken log₂ fold change, and the y-axis shows the −log₁₀ of the s-value. Orange dots denote genes passing both statistical significance and fold change thresholds; blue dots indicate statistically significant fold change without statistical significance in s-value; gray dots indicate no statistically significant fold change. c. Gene ontology (GO) groups of example upregulated (left) and downregulated (right) genes in the non-BaCl_2_-exposed worms that recently developed from tail pieces (Tail regenerate) versus non-BaCl_2_-exposed isolated worms (Control). Dot size refers to the number of associated genes, and the x-axis shows the −log₁₀(p-value) of each GO term.

### BaCl_2_ resistance activates bioelectric and immune responses distinct from canonical regeneration

Next, we sought to analyze the transcriptional signature of BaCl_2_ resistance. PCA of BaCl_2_-exposed worms and controls separates the groups (Figure 3a). This comparison revealed fewer differentially regulated genes compared to tail-regenerated: out of 39,721 genes, 69 (0.17%) were upregulated and 63 (0.16%) were downregulated (Figure 3b). Despite fewer overall changes, specific genes strongly indicated targeted bioelectric and immune responses. Among the top upregulated genes, the immune signaling regulator *traf3* (LFC=6.29) underscored robust innate immune activation in response to BaCl₂ exposure. Notably, the intracellular trafficking-related gene *KIAA1279* was significantly elevated again (LFC=4.80), indicating that altered intracellular transport is a shared aspect of both regeneration scenarios (Figure 3b). Downregulated genes in BaCl_2_-exposed worms included protease-related genes (*TMPRSS6*, LFC=– 1.73; *KLK13*, LFC=–1.65), mitochondrial dynamics regulators (e.g., Rapunzel proteins, LFC ∼–1.5), and lipid metabolism transporters (*NPC2*, LFC=–1.44) (Figure 3b).

**Figure 3.**
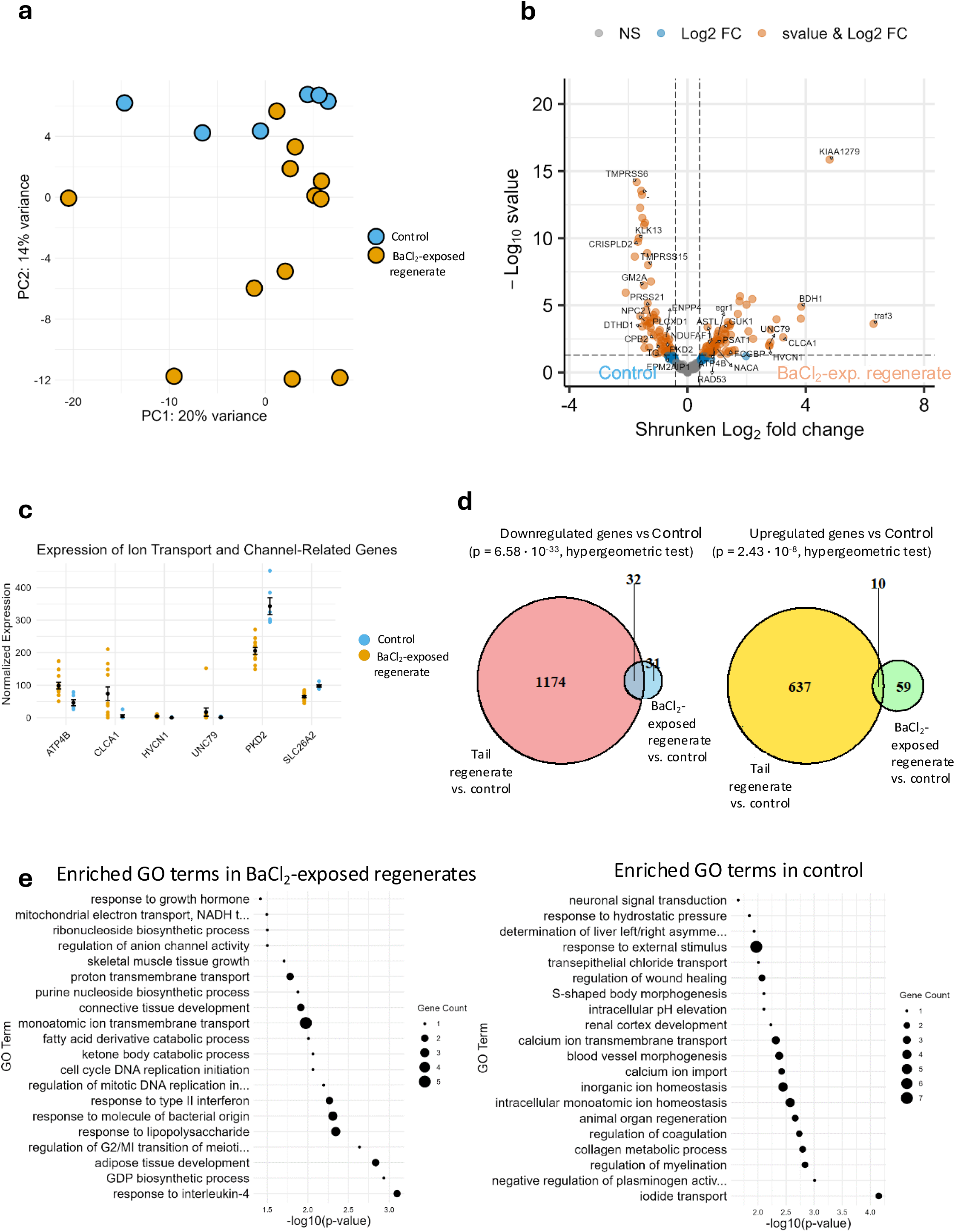
Transcriptional variation among BaCl_2_-exposed isolated regenerates (BaCl_2_ resistant) and non-BaCl_2_-exposed isolated worms (Control). a. Not all non-BaCl_2_-exposed isolated worms (Control: blue) and BaCl_2_-exposed isolated regenerates (BaCl_2_ resistant: orange) express the same levels of genes. PCA based on normalized gene expression from control and BaCl_2_-resistant samples. Each point represents a biological replicate. b. The volcano plot shows that 132 genes are differentially expressed in the BaCl_2_-exposed isolated regenerates (BaCl_2_ resistant) versus the non-BaCl_2_-exposed isolated worms (Control). The x-axis represents the shrunken log₂ fold change, and the y-axis shows the −log₁₀ of the s-value. Orange dots denote genes passing both statistical significance and fold change thresholds; blue dots indicate statistically significant fold change without statistical significance in s-value; gray dots indicate no statistically significant fold change. c. The normalized expression (y axis) of some ion-channel-related genes (x axis) shows that these genes are differentially expressed between BaCl_2_-resistant versus control worms. Each point represents a biological replicate colored by condition. d. 42 genes are differentially expressed in both the BaCl_2_-exposed isolated regenerates (B) and in the non-BaCl_2_-exposed worms that recently developed from tail pieces (Tail regenerate), in comparison to the non-BaCl_2_-exposed isolated worms (Control). Out of those 42 genes, 32 genes are both downregulated in both the BaCl_2_-exposed isolated regenerates (B) and in the non-BaCl_2_-exposed worms that recently developed from tail pieces (Tail regenerate), in comparison to the non-BaCl_2_-exposed isolated worms (Control). The remaining 10 genes are upregulated in both the BaCl_2_-exposed isolated regenerates (B) and in the non-BaCl_2_-exposed worms that recently developed from tail pieces (Tail regenerate), in comparison to the non-BaCl_2_-exposed isolated worms (C). P-values from hypergeometric tests indicate enrichment significance. e. Gene ontology (GO) groups of sample upregulated (left) and downregulated (right) genes in the BaCl_2_-exposed isolated regenerates versus non-BaCl_2_-exposed isolated worms excluding genes shared with the non-BaCl_2_-exposed worms that recently developed from tail pieces versus control comparison. Dot size represents the number of genes annotated with each term. The x-axis shows the –log₁₀(p-value).

Several genes directly involved in ion homeostasis and bioelectric regulation were prominently upregulated in the BaCl_2_-exposed worms. These included *CLCA1* (chloride channel regulator, LFC=3.23), *HVCN1* (voltage-gated proton channel, LFC=2.76), *UNC79* (NALCN sodium leak channel complex, LFC=2.75), and *ATP4B* (Na⁺/K⁺-ATPase subunit, LFC=0.90) (Figure 3c). These findings strongly suggest that a coordinated bioelectric adaptation underlies acquired resilience to BaCl_2_-induced ion channel blockade. These bioelectric adaptations may further influence tissue specification and regenerative outcomes by selectively modulating morphogenetic and ion transport programs during head reconstruction. Downregulated genes included planarian homologs of *PKD2* (LFC=–0.68, s-value=0.0049), a calcium-permeable cation channel involved in bioelectric signaling and membrane excitability, and *SLC26A2* (LFC=–0.54, s-value=0.0316), an anion transporter that regulates sulfate ion homeostasis and contributes to extracellular matrix organization. Taken together, these gene-level changes indicate a physiological reprogramming centered on ion transport resilience, immune defenses against microbes, and stress-buffering.

To determine whether the BaCl_2_-induced transcriptome recapitulates the program of canonical regeneration from tail fragments or represents a distinct transcriptional adaptation, we examined gene overlap between the two regeneration scenarios (Figure 3d). A modest yet statistically significant overlap existed. However, the majority of differentially expressed genes in BaCl_2_-exposed worms were unique. This indicates that BaCl_2_ adaptation constitutes a distinct response rather than a simple extension of the canonical regeneration program.

Functional enrichment analysis of BaCl₂-specific upregulated genes (i.e., excluding the regeneration-specific genes, Figure 2) revealed increased expression in pathways related to type II interferon response, response to bacterial molecules, adipose tissue development, skeletal muscle and connective tissue growth, regulation of anion channel activity, and proton and monoatomic ion transmembrane transport. Additional enriched categories included mitochondrial electron transport, purine biosynthesis, and hormonal signaling, suggesting a shift toward metabolic stabilization and immune readiness (Figure 3e).

Conversely, downregulated genes were enriched for myelination regulation, animal organ regeneration, calcium ion import, calcium ion transmembrane transport, intracellular pH elevation, S-shaped body morphogenesis, regulation of wound healing, transepithelial chloride transport, determination of left/right asymmetry, neural signal transduction, response to hydrostatic pressure, and iodide transport (Figure 3e).

### BaCl₂ Resistance Resolves into Two Transcriptional States

Planaria exhibit considerable individuality in behavior [59–61] and genetics [62]. PCA revealed a separation within the BaCl_2_-exposed group, forming two transcriptionally defined subclusters, B1 and B2, consisting of 10 and 2 worms, respectively (Figure 4a). Although the separation between these subclusters was statistically supported, the small size of B2 (n=2) limits confidence in defining it as a discrete transcriptional state; the distinction should therefore be considered suggestive rather than conclusive. Differential expression analysis between B1 and B2 revealed distinct transcriptional profiles (Figure 4b). The B2 subgroup, consisting of 2 worms, exhibited a set of genes strongly upregulated relative to group B1, including *CFAP43*, *ND4*, *SETMAR*, and *HIST1H3D*, suggestive of increased mitochondrial activity and chromatin remodeling. Conversely, B1 showed elevated expression of genes such as *SLC47A2*, *PKD2*, *UNC79*, and *RAG1*, pointing toward enhanced ion transport and immune-associated signaling (Figure 4b). Functional enrichment analysis further supported these divergent profiles. Genes upregulated in B1 (left panel, Figure 4c) were enriched for pathways involved in mitochondrial electron transport, proton transmembrane transport, oxidative and heavy metal stress responses (including copper and cadmium), and regulation of synaptic transmission and myelination. In contrast, genes upregulated in B2 (right panel, Figure 4c) were enriched for pathways related to immune modulation and cell fate regulation, including mucosal immune response, germ cell development, neurogenesis, vesicle docking, and negative regulation of potassium ion transport. These findings reveal that BaCl_2_ exposure response in *D. japonica* may emerge through just two distinct transcriptional programs; one dominated by bioelectric and oxidative stabilization mechanisms, and another centered on immune modulation and developmental reprogramming. These data are not consistent with a large number of possible solutions that could be individually found by each animal, but point to two attractors in the space of solutions.

**Figure 4.**
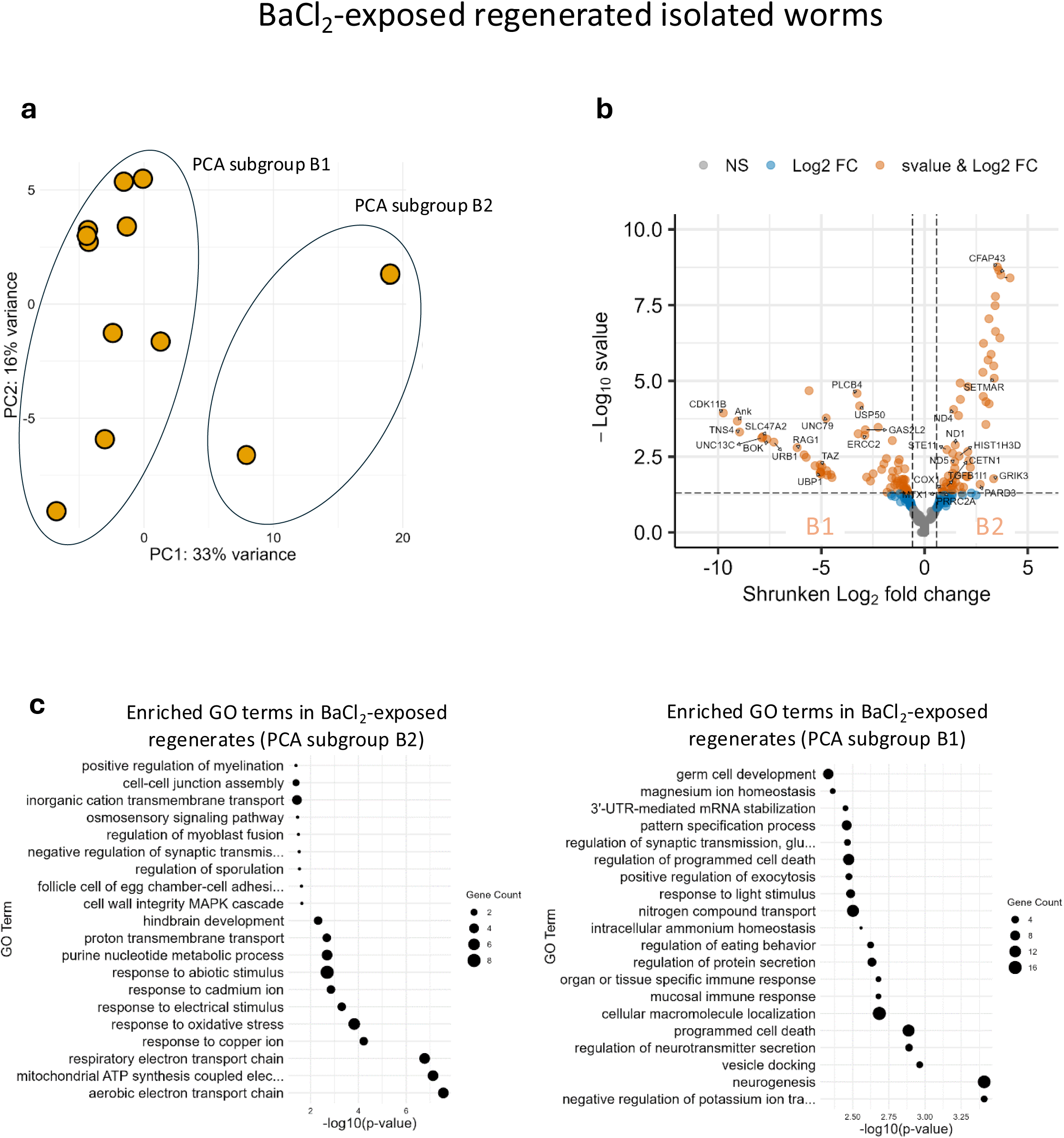
Two distinct transcriptional programs of BaCl_2_ adaptation in planaria. a. PCA plot based on normalized gene expression data from planarian samples. Each point represents a biological replicate. Transcriptional program B1: BaCl_2_-exposed isolated regenerates with some similarities in gene expression. Transcriptional program B2: BaCl_2_-exposed isolated regenerates with gene expression patterns not as similar to B1 worms. b. Volcano plot showing differentially expressed genes between putative BaCl_2_-resistant subgroups. The x-axis represents the shrunken log₂ fold change, and the y-axis shows the −log₁₀ of the s-value. Orange dots denote genes passing both statistical significance and fold change thresholds; blue dots indicate statistically significant fold change without statistical significance in s-value; gray dots indicate no statistically significant fold change. c. Gene ontology (GO) groups of sample upregulated (left) and downregulated (right) genes in B2 versus B1. Dot size corresponds to the number of associated genes, and the x-axis represents the −log₁₀(p-value) of each GO term.

### Isolated versus group-living worms in BaCl_2_ consistently differentially express two genes

To determine whether isolated *D. japonica* worms exhibit distinct transcriptomic profiles compared to those maintained in groups, we performed RNAseq on planaria that had regenerated their heads in BaCl_2_ while living in isolation versus together with 29 worms. We found that the isolated and group-living *D. japonica* worms do not differ in the majority of their transcripts (Figure S2b-e). Transcript dd_Djap_v4_66395_1_1 was upregulated in all comparisons of isolates versus each replicate of the group-living worms (Figure S2c-e; Table S1). A homolog of this gene plays a role in regulating the activity of choline O-acetyltransferase. Another transcript (dd_Djap_v4_56151_2_1) was downregulated in isolated versus group-living worms (Figure S2c-e; Table S1). A homolog of this gene encodes for the enzyme leishmanolysin. Under BaCl₂ exposure, social isolation produced only minimal transcriptomic differences, with two transcripts consistently differing between conditions. These results suggest that cohort size has limited impact on the global transcriptional response in head-regenerated worms. The small number of differentially expressed transcripts precludes broader functional interpretation.

## Discussion

Our transcriptomic analysis reveals that *D. japonica* employs distinct molecular programs depending on the regenerative context, with head regeneration under BaCl_2_ stress diverging from the canonical regeneration of head from tail pieces. We observe that recent regenerative history has a stronger effect on global gene expression than exposure to BaCl₂ alone, however, no recent regeneration and regeneration in BaCl₂ may involve more transcriptomic flexibility than canonical anterior regeneration. Note that both barium-treated and untreated regenerating worms have to rebuild their heads. The fact that BaCl₂-treated worms do so without involving most of the generic regeneration response genes suggests the possibility that the process of finding a BaCl₂-compatible gene expression profile also allowed them to avoid expression default regeneration genes that are not actually necessary to produce a functioning head. It is also possible however that the generic genes were indeed activated but with a novel timing that allowed them to escape detection by our sampling strategy.

While both regeneration scenarios share common transcriptional features like intracellular (e.g., upregulation of *KIAA1279*) and stress management, they diverge in their dominant biological strategies. Tail-regenerated worms are characterized by robust activation of developmental, cytoskeletal, and oxidative pathways, supporting structural rebuilding. In contrast, response to BaCl_2_-exposure prioritizes bioelectric homeostasis, immune activation, and selective metabolic adaptation, reflecting a distinct transcriptional solution to an environmental ion channel blockade. In this dataset, BaCl₂-exposed head-regenerating worms segregated into one dominant transcriptional cluster and a small secondary cluster. While this pattern indicates structured heterogeneity within the population, much larger sample sizes in future work could better delineate the possible number of distinct solutions that might be available overall.

### Selective upregulation of head morphogenesis programs in tail-regenerated worms

Recent tail-regenerated worms living in isolation downregulate a broad set of genes while selectively upregulating head-related morphogenesis genes. This could reflect an energy-conserving strategy [63, 64] that prioritizes head reconstruction over general maintenance. These findings align with previous studies in *Schmidtea mediterranea* and reinforce the notion that regeneration induces long-lasting molecular changes. The upregulation of genes related to morphogenesis of head structures in the tail-regenerated worms is similar to previous findings of tail-regenerated *S. mediterranea* [65].

### Differential regulation of mitochondrial quality control pathways in controls

Comparative transcriptomic analysis revealed enrichment of genes associated with mitochondrial depolarization, mitophagy, and mitochondrial fusion in controls relative to tail-regenerated worms. Given that mitochondrial depolarization is a known precursor to mitophagic clearance [66–68], these findings are consistent with active mitochondrial turnover. Similar enrichment of autophagy-related pathways has been reported in *D. japonica* tissues [69]. While functional validation is required, the data are compatible with differential regulation of mitochondrial quality control processes across groups. Mitophagy may be an additional component, together with other known planarian mechanisms (e.g., telomere maintenance, efficient DNA repair, silencing of transposable elements) [70], contributing to planarian immortality.

### BaCl_2_ exposure affects flatworm bioelectrics and the flatworm-microbial interactome

Compared to prior studies focusing on head tissue [6], our whole-worm transcriptomic analysis— controlled for regeneration effects—reveals downregulation of chloride and calcium transport pathways, likely mitigating ionic imbalance caused by BaCl₂. Emmons-Bell et al. [6] had seen an upregulation of an ion channel called TRPMa in BaCl_2_-exposed heads suggesting that these additional TRPMa channels may be helping the planaria adapt in BaCl_2_. However, Emmons-Bell et al. [6] did not control for transcriptomic changes that could be due to regeneration. The upregulation of the beta subunit of the gastric H^+^/K^+^ ATPase pump in BaCl_2_-exposed worms (barium non-specifically blocks potassium channels) is related to previous findings in rabbits that in the absence of K^+^ the H^+^/K^+^ ATPase pump is linked with the barium-sensitive pathway [71], and consistent with prior data showing the H^+^/K^+^ ATPase pump being involved in planarian regenerative polarity [72].

In this study, we were able to control for regeneration-related transcriptomic changes due to our experimental design including tail-regenerated worms in the absence of BaCl_2_. Thus, by excluding the differentially expressed genes that were related to regeneration and using whole worms in our transcriptomic analysis, we also found downregulation of iodide transport and downregulation of transepithelial chloride transport, similar to the downregulation of innexin [6]. The latter observation reinforces that worms are limiting the spread of chloride ions to other cells, thus limiting the effect of BaCl_2_ on their tissues, and enabling planarian survival and head regeneration in BaCl_2_. Previous studies in sympathetic neurons of chick embryos and murine Bal17 B cell lymphoma cells have shown that Ba^+2^ prevents Ca^+2^ from exiting the cell and triggers the release of Ca^+2^ from intracellular locations to the cytoplasm [73, 74]. Therefore, the downregulation of genes related to calcium import and calcium transmembrane transport in BaCl_2_-exposed worms may be a response of the worms to minimize the accumulation of Ca^+2^ in the cytoplasm.

Concurrent upregulation of immune genes may reflect microbial proliferation in the medium due to head degradation. Our analysis also showed upregulation of many genes related to immunity against microbes. Williams et al. had found that bacteria, specifically the endogenous commensal *Aquitalea* sp. bacteria of *D. japonica* planaria, not only affect the regeneration of planaria, such as how many heads or eyes they develop, but also the patterning of the planarian nervous system [75]. Therefore, BaCl_2_ may be altering the interaction between bacteria and the worms’ nervous system. Also, debris from the degraded planarian head may be allowing increased bacterial growth in the medium and thus the upregulation of antimicrobial genes in the planaria may be a response against an increased concentration of microbes in the medium. Concurrent upregulation of immune genes may reflect microbial proliferation in the medium due to head degradation

### Regeneration under BaCl₂ exposure occurs largely independently of social context

Isolated planaria regenerated heads under BaCl₂ exposure and exhibited transcriptional states comparable to those observed in group-housed worms. This indicates that successful regeneration under ionic stress does not require communal signaling or social buffering. The presence of similar transcriptional programs in independently living worms suggests that successful regenerative response to BaCl₂ may be intrinsically encoded rather than socially coordinated. However, it is also possible that group dynamics occur at levels other than transcriptional, and would require other assays to be observed.

### Limitations of our study

Transcript-level expression was quantified using a reference transcriptome for *D. japonica*, followed by aggregation to gene-level counts using a functionally annotated transcript-to-gene mapping (see Methods). While this enabled robust gene-level quantification across samples, many transcripts remained unmapped or unannotated, reflecting the incomplete characterization of the *D. japonica* transcriptome. Another fundamental limitation, of the fact that transcriptomics is destructive, is that it is not yet possible to characterize the time-dependent transcriptional course of the same worm in transcriptional space. Future advances in technology will allow determination of whether the *paths* of different worms toward the same solution to BaCl_2_ challenge are the same or differ.

## Conclusion

In conclusion, planaria possess a remarkable capacity for transcriptional problem-solving in response to physiological stress and regenerative demands. Tail-regenerated worms prioritize morphogenesis, and downregulate mitochondrial recycling, suggesting distinct strategies for tissue renewal and longevity. Exposure to BaCl₂ elicits at least two divergent transcriptional programs, highlighting variability in stress adaptation. These insights are a stepping-stone for future studies investigating the exact mechanism by which planaria find one transcriptomic solution to the BaCl_2_ challenge in the absence of others, and the social structure of chemical resilience in other organisms beyond planaria. The results underscore the autonomous and flexible nature of planarian regeneration and resilience, positioning this model organism as a powerful system for exploring the molecular foundations of adaptation, longevity, and environmental response.

## Data and code availability

The data and code used in the manuscript are available in NCBI (GEO Accession number GSE341203; https://www.ncbi.nlm.nih.gov/geo/query/acc.cgi?acc=GSE341203).

## Acknowledgements

We thank Angela Tung for advice in performing the barium chloride experiments, Vaibhav Pai for advice on extracting RNA from individual planaria, Albert Tai, Rakela Colon, and Jayati Mandal for helping arrange the process of RNA sequencing, and Robert Brucker for many helpful comments on the manuscript.

## Author contributions

**S.E.K.:** Data curation, software, formal analysis, investigation, methodology, writing–original draft, writing–review and editing, performed the experiments. **T.L.:** Data curation, software, formal analysis, investigation, methodology, writing–original draft, writing–review and editing. **M.L.:** Conceptualization, supervision, funding acquisition, formal analysis, investigation, methodology, project administration, writing–review and editing.

## Supplementary figure legends

<u>Figure S1. Read counts from the transcriptomes of each worm vary within and among the different conditions.</u> Non-BaCl_2_-exposed isolated worms (Control); non-BaCl_2_-exposed worms that recently developed from tail pieces (Tail regenerate); BaCl_2_-exposed isolated regenerates (BaCl_2_–exposed regenerate). Each number on the x axis represents a different worm. B8, B8.2, and B8.3 are samples from the same worm.

<u>Figure S2. BaCl**_2_**-exposed regenerated worms living on their own versus in groups do not differ statistically significantly in their transcriptome, apart from two genes.</u>

a. Design of experiment testing for interindividual differences in the transcriptome of non-BaCl_2_-exposed non-fissioned planaria, non-<u>BaCl_2_</u>-exposed planaria that recently developed from tail pieces of fissioned planaria, and regenerated planaria that had lost their head in BaCl_2_.

b. Group-living (G) and isolated (S) BaCl_2_-exposed regenerates show minimal transcriptomic differences. G: group-living BaCl_2_-exposed regenerates. S: isolated BaCl_2_-exposed regenerates. a. G1: 4 worms, G2: 4 worms, G3: 4 worms; G: 12 worms; S: 3 worms. c-e. The genes in red with a positive log_2_ fold change are overexpressed in isolated versus group-living worms, whereas the genes in red with a negative log_2_ fold change are underexpressed in isolated versus group-living worms.

c. Comparison of gene expression levels in group living worms (4 worms living in a single Petri dish: G1) versus isolated worms (3 isolated worms, each living in a different Petri dish). The x-axis represents the shrunken log₂ fold change, and the y-axis shows the −log₁₀ of the s-value. Red dots denote genes passing both statistical significance and fold change thresholds; green dots indicate statistically significant fold change without statistical significance in s-value; gray dots indicate no statistically significant fold change.

d. Comparison of gene expression levels in group living worms (4 worms living in a single Petri dish: G2) versus isolated worms (3 isolated worms, each living in a different Petri dish). e. Comparison of gene expression levels in group living worms (4 worms living in a single Petri dish: G3) versus isolated worms (3 isolated worms, each living in a different Petri dish). There are only two differentially expressed genes between the group-living (G) versus isolated (S) worms that lost and then regenerated their heads in BaCl_2_ (c-e). Large red circles highlight the genes that are statistically differentially expressed in groups-living versus isolated worms.

<u>Table S1. Functional and organismal information of two differentially expressed genes in isolated versus group-living planaria that lost and regenerated their heads in BaCl_2_.</u>

