## Supplementary material for "Transcriptional Profiling of Planarian Regeneration Habituating to Physiological Stressor Reveals Individual and Collective Dynamics": Table S1

| **Differentially expressed gene information** | **transcript_id** | **gene_id** | **gene_symbol** | **organism** | **GOs** | **evalue** | **Description** | **KEGG_Pathway** |
| --- | --- | --- | --- | --- | --- | --- | --- | --- |
| This gene is **upregulated** in singleton barium-treated worms in all S-G1, S-G2, S-G3 comparisons. | dd_Djap_v4_66395_1_1 | 6183.Smp_167120.1 | - | 32208\|Metazoa | - | 1.47 x 10^-30^ | regulation of choline O-acetyltransferase activity | - |
| This gene is **downregulated** in singleton barium-treated worms in all S-G1, S-G2, S-G3 comparisons. | dd_Djap_v4_56151_2_1 | 65489.OBART03G33000.2 | - | 35493\|Streptophyta | 3.4.24.36 | 6.28 x 10^-17^ | Leishmanolysin | ko05140, ko05143, map05140,  map05143 |
