## Supplementary Figures 1 & 2 for "Transcriptional Profiling of Planarian Regeneration Habituating to Physiological Stressor Reveals Individual and Collective Dynamics"

### Slide 1
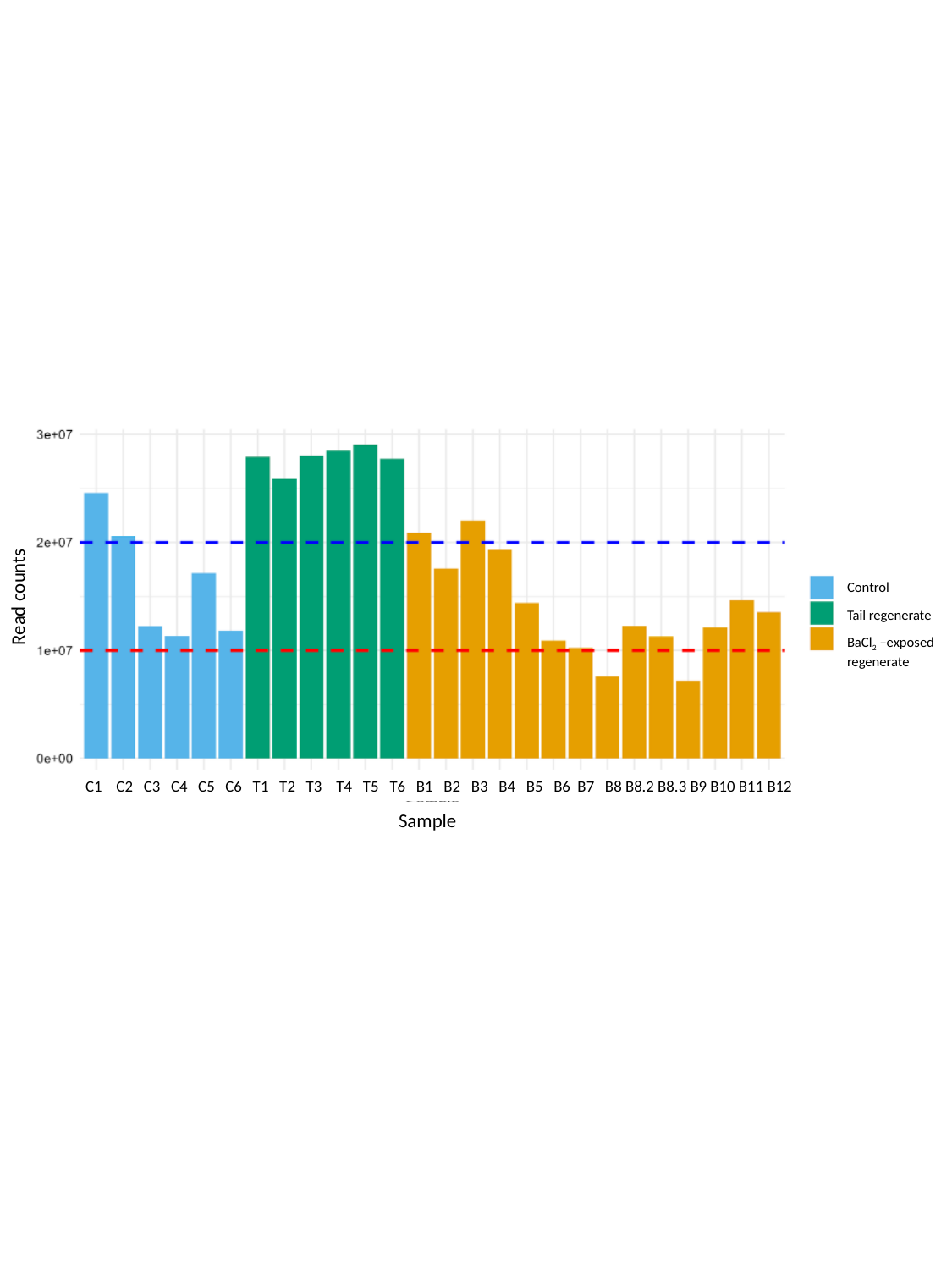

Control
Tail regenerate
BaCl_2_ –exposed regenerate
Read counts
C1 C2 C3 C4 C5 C6 T1 T2 T3 T4 T5 T6 B1 B2 B3 B4 B5 B6 B7 B8 B8.2 B8.3 B9 B10 B11 B12
Sample

### Slide 2
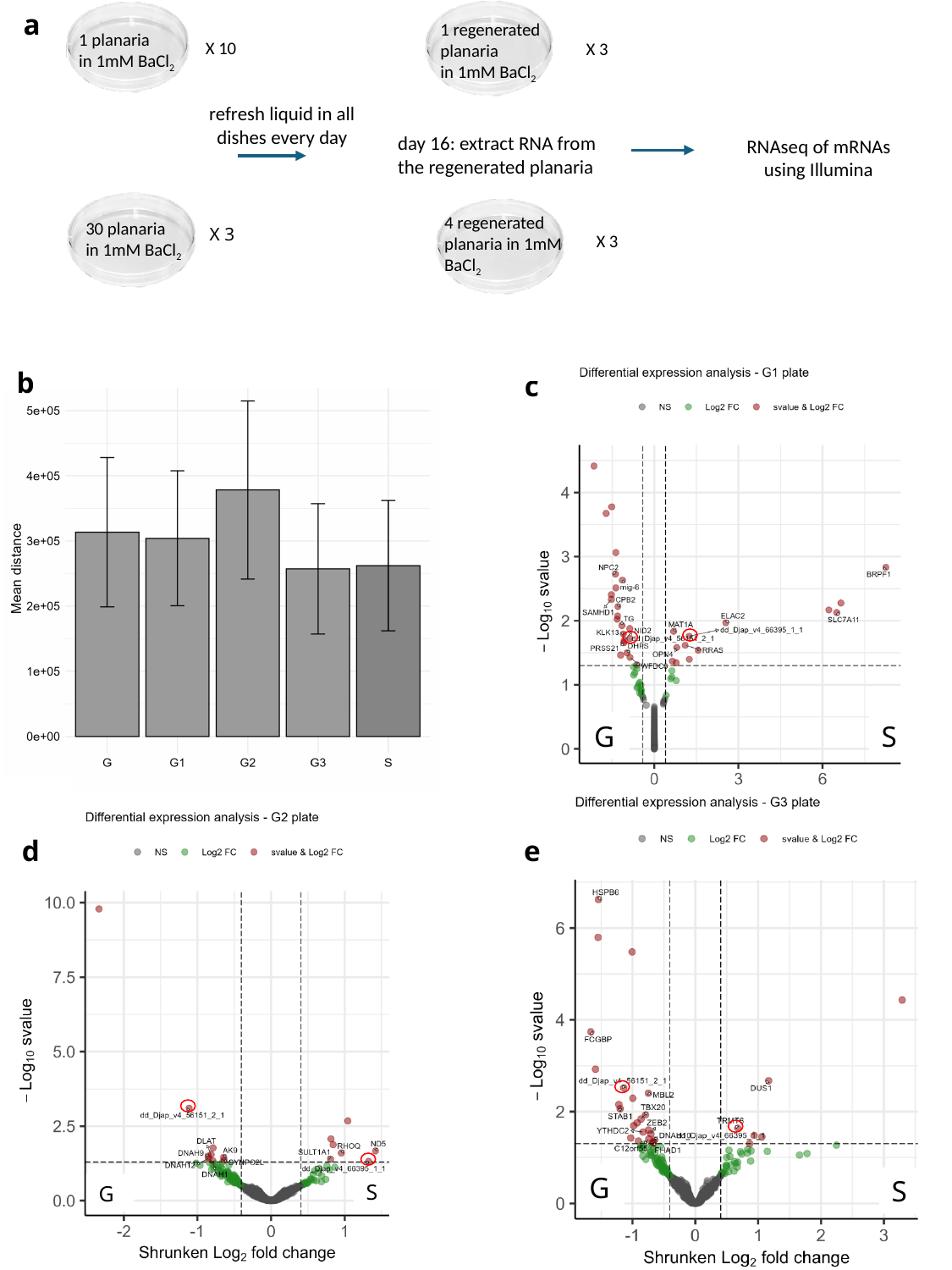

a
1 regenerated planaria
in 1mM BaCl_2_
1 planaria
in 1mM BaCl_2_
X 10
X 3
refresh liquid in all dishes every day
day 16: extract RNA from the regenerated planaria
RNAseq of mRNAs using Illumina
4 regenerated planaria in 1mM BaCl_2_
30 planaria
in 1mM BaCl_2_
X 3
X 3
b
c
G
S
e
d
G
S
S
G
